# Optogenetic currents in myofibroblasts acutely alter electrophysiology and conduction of co-cultured cardiomyocytes

**DOI:** 10.1101/2020.06.02.124529

**Authors:** Geran Kostecki, Yu Shi, Christopher Chen, Daniel H. Reich, Emilia Entcheva, Leslie Tung

**Affiliations:** Department of Biomedical Engineering, Johns Hopkins University, Baltimore, MD, USA; Department of Physics and Astronomy, Johns Hopkins University, Baltimore, MD 21218, USA; Biological Design Center, Department of Biomedical Engineering, Boston University, Boston, MA; Wyss Institute for Biologically Inspired Engineering, Harvard University, Boston, MA; Department of Biomedical Engineering, George Washington University, Washington, DC USA

## Abstract

Interactions between cardiac myofibroblasts and myocytes may slow conduction after cardiac injury, increasing the chance of life-threatening arrhythmia. While co-culture studies have shown that myofibroblasts can affect cardiomyocyte electrophysiology *in vitro*, the mechanism(s) remain debatable. In this study, primary neonatal rat cardiac myofibroblasts were transduced with the light-activated ion channel Channelrhodopsin-2, which allowed acute and selective modulation of myofibroblast currents in co-cultures with cardiomyocytes. Optical mapping revealed that myofibroblast-specific optogenetically induced inward currents decreased conduction velocity in the co-cultures by 27±6% (baseline = 17.7±5.3 cm/s), and shortened the cardiac action potential duration by 14±7% (baseline = 161±11 ms) when 0.017 mW/mm^2^ light was applied. When light irradiance was increased to 0.057 mW/mm^2^, the myofibroblast currents led to spontaneous beating in 6/7 co-cultures. Experiments showed that optogenetic perturbation did not lead to changes in myofibroblast strain and force generation, suggesting purely electrical effects in this model. *In silico* modeling of optogenetically modified myofibroblast-cardiomyocyte co-cultures largely reproduced these results and enabled a comprehensive study of relevant parameters. These results clearly demonstrate that myofibroblasts are sufficiently electrically connected to cardiomyocytes to effectively alter macroscopic electrophysiological properties in this model of cardiac tissue.

## Introduction

After myocardial injury, paracrine factors, including transforming growth factor-β1 (TGF-β1), cause fibroblasts to differentiate into myofibroblasts (MFBs) [1–5]. MFBs express α-smooth muscle actin (α-SMA) fibers and contract, which stabilizes and shrinks the injured area. They also secrete extracellular matrix which replaces dead cells and further mechanically stabilizes the tissue [1–5].

Along with these changes, arrhythmia risk is significantly increased [6, 7]. There are many contributing factors to this, such as increased heterogeneity of cardiomyocyte (CM) coupling within the scar causing zigzag propagation or electrical block, as well as ion channel remodeling in the CMs themselves [5–8]. However, an additional factor may be the effects of the MFBs on CM electrophysiology, since they can remain in a differentiated state in the injured area years after injury [9]. Simplified *in vitro* systems have shown increased occurrence [10] and complexity [11] of spiral waves with increasing fraction of MFBs in co-culture. This is likely because addition of MFBs to CMs slows CM conduction velocity (CV) [11–18], and increases spontaneous beating rate [14, 15, 18, 19]. It is commonly suggested that such effects are due to electrical coupling between CMs and the less electrically polarized MFBs which causes current to flow into CMs at rest [14, 16, 18], thereby raising CM resting potential and inactivating sodium channels or generating spontaneous beating [11, 14, 16–18]. However, these pro-arrhythmic MFB-CM interactions may instead be caused by paracrine [8, 20, 21] and mechanical [12, 13, 15] mechanisms.

In this study, MFBs were transduced with Channelrhodopsin-2 (ChR2), a relatively non-selective cation channel that opens in response to light [22], to acutely depolarize them. This MFB-specific perturbation enabled the study of acute effects of MFB depolarizing currents on macroscopic electrophysiological properties in syncytia of co-cultured MFBs and CMs.

## Methods

### Cell culture

Neonatal rat ventricular CMs were produced as described previously [23], with minor modifications, and plated onto coverslips coated with 25 μg/mL fibronectin at 1 million cells per well for a 12-well plate or 500,000 per well for a 24-well plate (approximately 250,000/cm^2^). During isolation, CMs were separated from fibroblasts using two 1-hour preplating steps. Fibroblasts from the first preplate were passaged twice. Some were transduced during the second passage with ChR2-YFP adenovirus at a multiplicity of infection of 2,000, as determined by experiments (Supp. Fig. 1), with media changed 4-6 hours later to remove virus as described previously [24]. Transduced and untransduced fibroblasts were treated with 5 ng/mL TGF-β1 (R+D Systems) for 2-3 days to differentiate them into MFBs.

### Co-culture, imaging, and optical mapping

ChR2-transduced MFBs (ChR2-MFBs) or untransduced MFBs were added to 4-5-day-old CM monolayers at 400,000/well in 12-well plates or 200,000/well in 24-well plates (approximately 100,000/cm^2^), giving a MFB:CM cell ratio of 0.4. High levels of TGF-β1 treatment were used to irreversibly differentiate fibroblasts into MFBs [25] to ensure maintenance of an MFB phenotype without application of exogenous TGF-β1 during co-culture, which could have directly affected CMs. On days 5-8, co-cultures were imaged under phase contrast and fluorescence (Eclipse TE2000U, Nikon) to examine their morphology and continued expression of ChR2, and then placed in a custom optical mapping system [26] and stained for 5 minutes with 35 μM of the voltage-sensitive dye di-4-ANBDQBS (obtained from Dr. Leslie Loew, University of Connecticut), which is excited by red light (λ = 655 nm) [27] and therefore spectrally compatible with ChR2 [22, 28, 29]. Tyrode’s solution at 35°C was then continuously flowed over the cells. The pacing threshold was determined to within 1 V, and the cells were point paced at 1.1x threshold for 5 min at 500 ms cycle length (CL) to reach steady state. A baseline optical recording was taken, and then continuous blue light (λ = 448 nm) was applied across the entire monolayer to activate the ChR2 channels for approximately 2 s before and throughout the duration of a 2 s recording, after which the light was switched off, and a post-activation recording was collected within seconds. This was done for different light intensities, starting at approximately the lowest intensities for which changes could be detected (I_0_ = 0.0057 mW/mm^2^), and increasing to 3*I_0_ and 10*I_0_, at which point most samples beat spontaneously faster than the paced rate. Co-cultures were fixed and stained for α-actinin (Sigma), α-SMA (DAKO), connexin43 (Cx43) (Sigma), YFP (Invitrogen GFP Ab), and/or DAPI before confocal imaging (LSM 710NLO-Meta, Zeiss), and plate fluorescence reader measurements (Spectramax M2, Molecular Devices).

### Mathematical model

Experimental data were modeled in MATLAB (The MathWorks) using a modified version of the Korhonen model of neonatal rat ventricular CMs [30] (see Online Data Supplement for modification details). CMs were connected via a lumped gap junction to a lumped MFB compartment, with currents assumed to be time-independent and modeled by a hyperbolic fit to the I/V curve measured by Salvarani et al. for MFBs [14] (Fig. 1A). In addition to endogenous currents, the Williams model for ChR2 current [31] was added to MFBs. MFBs were considered to have the same capacitance per cell as CMs, as done previously [32], and so the MFB:CM capacitance ratio was 0.4 to match the cell number ratio in experiments. Each CM was 50 μm long, and had a 50 µm-long interface with the following cell, similar to the cell length found by Jousset [32]. A 30-cell-long 1-D strand of ChR2-MFB/CM cell pairs was modeled (Fig. 1B). Based on previous data which found conductance between two CMs to be 7.7 nS/μm [14], neighboring CMs were connected by lumped gap junctions with a conductance of 50 μm*7.7 nS/μm = 385 nS. Neighboring MFBs were not connected to each other to prevent them from having “double-sided” effects on CMs [33] and creating an alternate current path. Because the MFB-CM coupling conductance (G_MFB-CM_) is unknown for MFBs lying on top of CMs, and because ChR2 conductance (g_ChR2_) varies considerably depending on cell type [31], maximum likelihood estimation was used to determine which values best fit our experimental ΔCL, ΔCV, and ΔAPD data. These values were then used to compare our model to the data. The effects of a wider range of G_MFB-CM_ (10^−1^-10^2^ nS/CM) and light intensities (0.3*I_0_-300*I_0_) on spontaneous beating and CV was also modelled.

**Figure 1.**
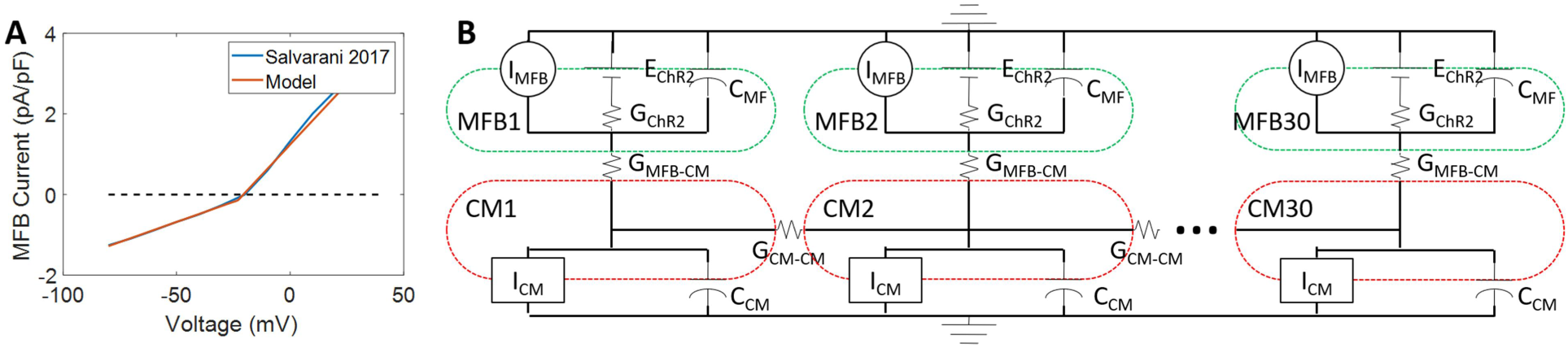
Computational model. **A.** Endogenous cardiac myofibroblast (MFB) currents were modeled by fitting a hyperbola to steady-state current-voltage measurements in TGF-β-treated MFBs measured by Salvarani, et. al. **B.** The ChR2 channel model (G_ChR2_, E_ChR2_) from Williams, et. al. was added to the endogenous currents. MFBs were electrically connected (G_MFB-CM_) to neonatal rat cardiomyocytes (CMs) as modeled by Korhonen, et. al. which were connected to each other (G_CM-CM_) to form a 30-cell, 1.5 mm 1-D cable. The dashed lines show the outer boundaries of the CMs (red) and MFBs (green).

### Data processing and statistics

Optical mapping data were processed by custom MATLAB software. Co-cultures with initial CV below 10 cm/s were excluded from analysis. Confocal images were processed by FIJI [34] and Zen Black (Zeiss) software. Background values from empty wells were subtracted from plate reader measurements. All data are presented as mean±SD. Paired or unpaired t-tests with unequal variances were used to determine statistical differences, where appropriate. The t-distribution was used for maximum likelihood estimation. Differences were considered statistically significant at p<0.05.

## Results

A substantial portion of this work has been previously reported in a Ph.D. thesis [35].

### Co-culture of cardiomyocytes with ChR2-transduced myofibroblasts

To assess whether inward current in MFBs can affect CM electrophysiology, fibroblasts were transduced with ChR2 and differentiated into MFBs by treatment with TGF-β1, then co-cultured with CMs. Confocal imaging of MFBs with CMs demonstrated continued expression of α-SMA by MFBs 2 days after plating on CMs and concomitant cessation of TGF-β1 treatment (Fig. 2A-C). Wide field (∼2 mm) imaging of co-cultures showed the plated ChR2-MFBs formed a homogeneous, dense network over a wide area of CMs (Fig. 2D) and continued to express ChR2 during co-culture (Fig. 2E). Confocal imaging showed confluent CMs (Fig. 2F) with ChR2-MFBs resting on top of them (Fig. 2G and H), as well as Cx43 puncta in the MFB cell layer, suggesting expression of Cx43 by MFBs (Fig. 2G).

**Figure 2.**
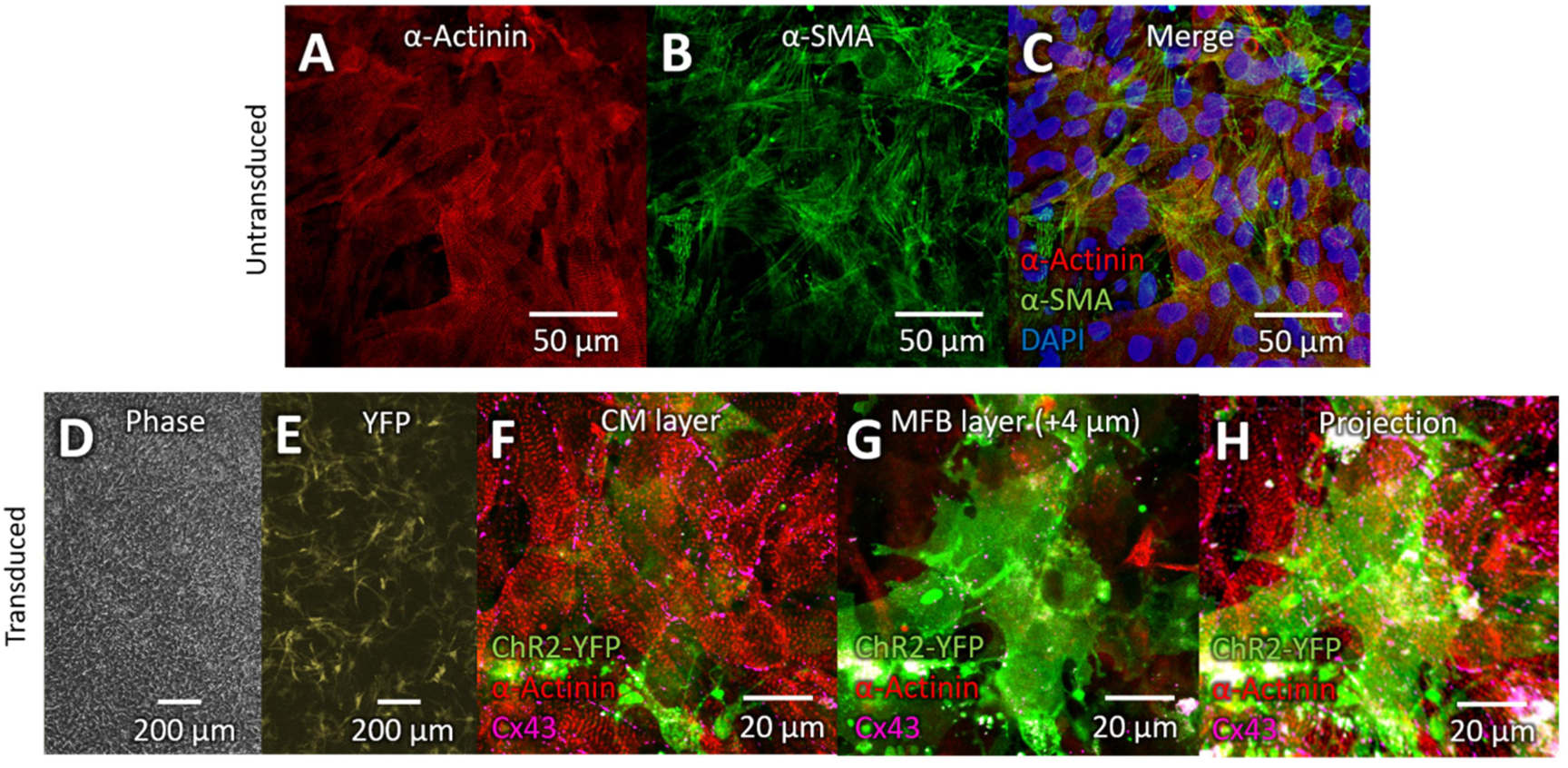
Co-culture of cardiomyocytes and ChR2-transduced myofibroblasts. ***A-C.*** Confocal image of co-culture of MFBs and CMs. α-actinin (**A**, red) marks CMs, and α-smooth muscle actin (**B**, α-SMA, green) marks MFBs. **C.** Merge of **A** and **B** with DAPI (blue) to stain nuclei. **D.** Phase contrast image of CM co-cultured with ChR2-MFBs. **E.** Fluorescence image of same sample as **D**, with YFP marking transduced MFBs. **F-H.** Confocal image of transduced MFBs and CMs. ChR2-YFP (green) marks transduced MFBs, α-actinin (red) marks CMs, and violet shows connexin43 (Cx43). **F.** CM layer of Z-stack showing gap junctions between CMs. **G.** Image from 4 μm above F showing MFBs on top of CMs, as well as Cx43 puncta, apparently between CMs and MFBs. **H.** Maximum intensity projection of entire 18 μm-thick z-stack sampled every 2 μm.

### Myofibroblast currents can cause electrophysiological changes in cardiomyocytes

In ChR2-MFB/CM co-cultures electrically paced at 500 ms CL (Fig. 3Ai), application of continuous blue light opened ChR2 channels and could induce spontaneous beating at a rate higher than the 500 ms paced CL (Fig. 3Aii). Cessation of light (and therefore ChR2 current) caused spontaneous beating to stop (Fig. 3Aiii). This spontaneous beating in response to light occurred in almost all ChR2-MFB/CM co-cultures, but in none of the MFB/CM co-cultures (Fig. 3B), thus precluding thermal or other non-specific effects of the applied light. Data across multiple samples showed that 3*I_0_ light caused spontaneous beating faster than the 500 ms paced CL in 4/10 samples, and that the beating CL decreased as the ChR2 current increased (Fig. 3B). CL of ChR2-MFB/CM co-cultures under 10*I_0_ light was reduced significantly reduced (ΔCL = −36±18% from a baseline of 500±1 ms, n=7, p=0.002, Fig. 3B).

**Figure 3.**
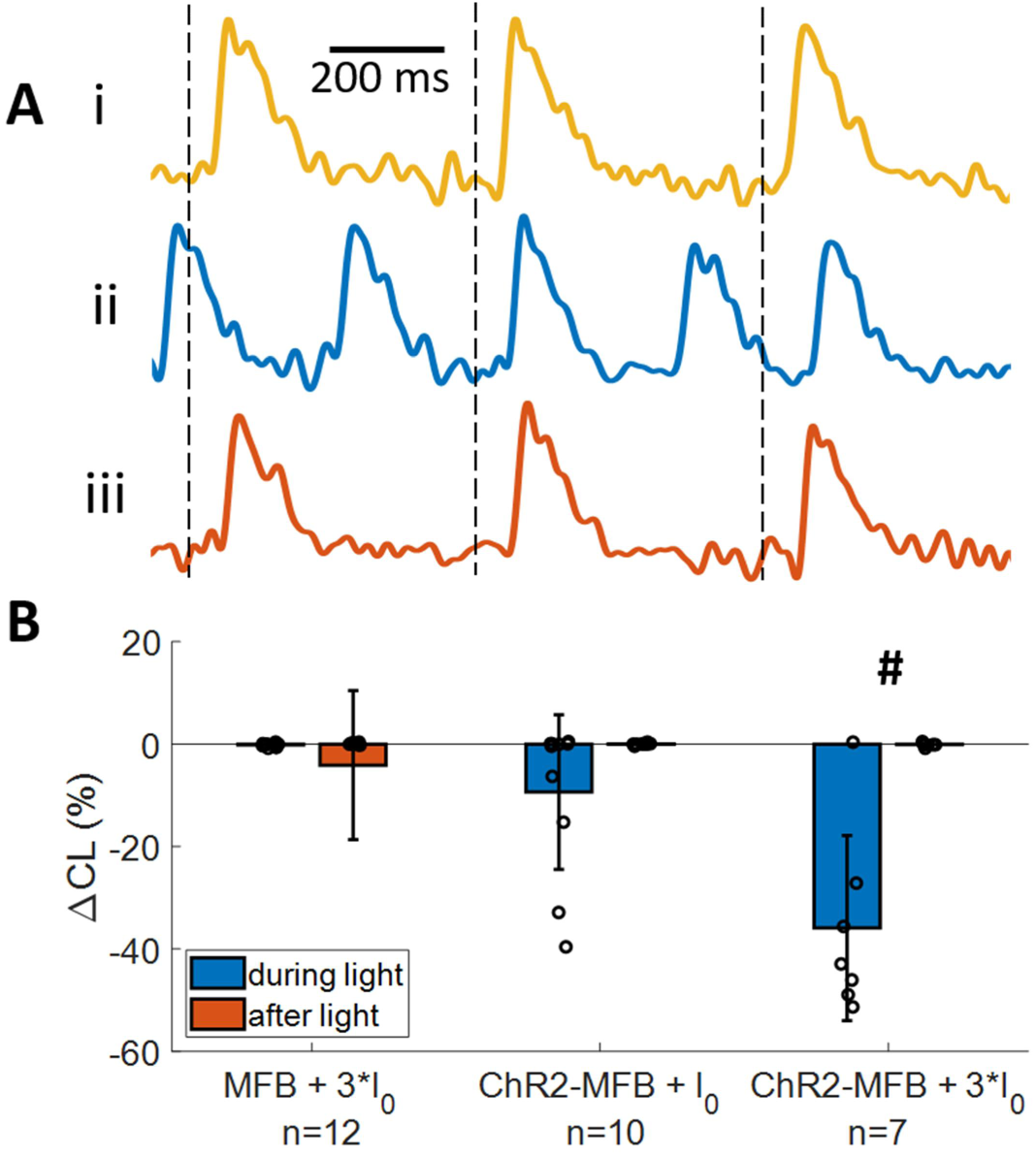
Inward currents in myofibroblasts cause spontaneous beating in co-cultured cardiomyocyte syncytia. **A.** Voltage traces of a co-culture of ChR2-transduced MFBs (ChR2-MFBs) with CMs before (I, gold), during (ii, blue), and after (iii, orange) application of 3*I_0_ blue light (I_0_ = 0.0057 mW/mm^2^, approximately the lowest light intensity at which functional effects were observed) to activate ChR2 current in MFBs. Dashed lines show time of pacing. **B.** Percent change in cycle length (CL) (vs. before light application) during and after application of light at different power during 500 ms CL pacing, for co-cultures of CM with MFBs or ChR2-MFBs. # shows p<0.005 difference between ChR2-MFB/CM and MFB/CM co-cultures.

Addition of ChR2-MFBs trended towards reducing CV relative to pure CM cultures during 500 ms CL pacing (17.7±5.3, n=8 vs. 20.9±4.3, n=14, respectively, p=0.17). For ChR2-MFB/CM co-cultures (Fig. 4Ai), application of light slowed CV (Fig. 4Aii). This slowing was reversed when the light was turned off (Fig. 4Aiii). CV decreased further as light intensity was increased, until the onset of spontaneous beating prevented further comparison of CVs (since CV can change from beating rate changes alone) (Fig. 4B). Slowing occurred specifically in ChR2-MFB/CM co-cultures in a dose-dependent manner across multiple samples (Fig. 4B). There was significant slowing in ChR2-MFB/CM co-cultures at light levels as low as I_0_ (ΔCV = −12±11% from a baseline of 17.7±5.3 cm/s, n=8, p=0.01). Slowing was exacerbated at 3*I_0_ (ΔCV = −27±5% from a baseline of 14.8±3.6 cm/s, n=5, p<10^−3^, Fig. 4B).

**Figure 4.**
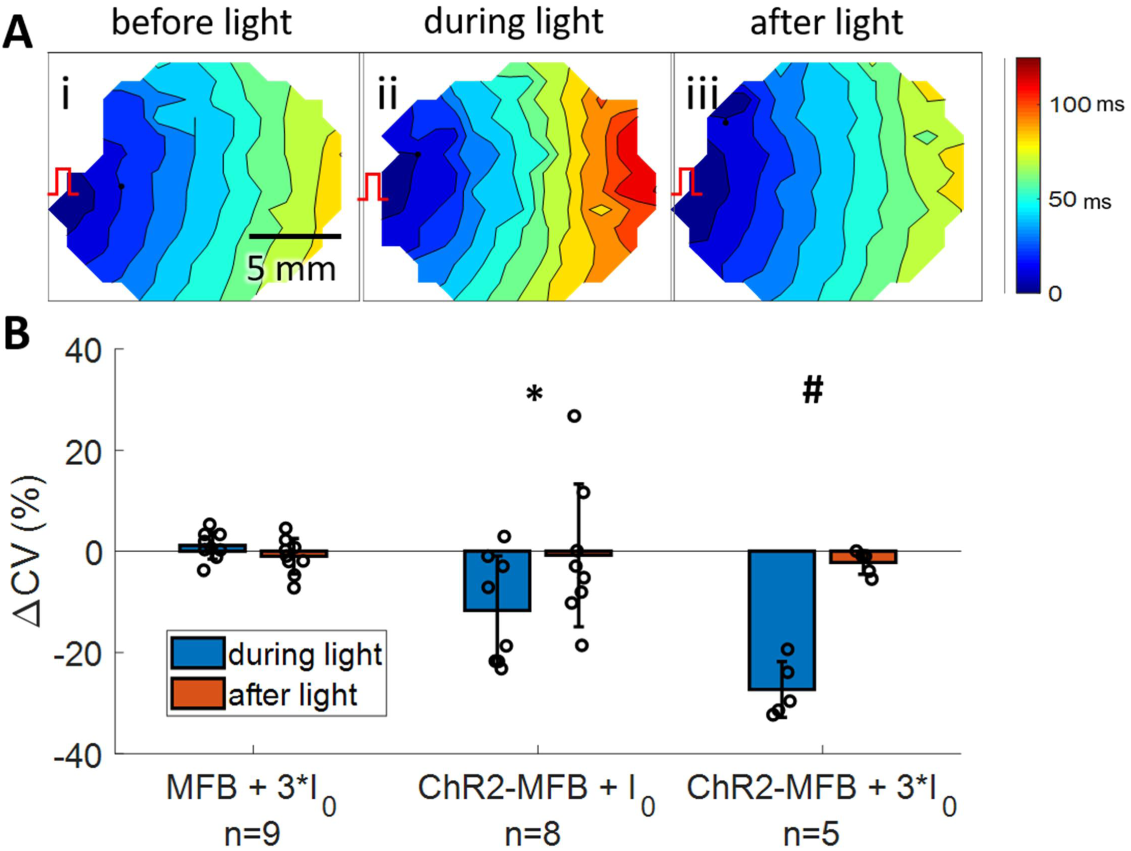
Inward currents in myofibroblasts cause slowing in co-cultured cardiomyocyte syncytia. **A.** Activation maps of a co-culture of ChR2-MFBs with CMs before (**i**), during (**ii**), and after (**iii**) application of 3*I_0_ blue light to activate ChR2 current in MFBs. Color bar at right shows activation time scale. Isochrones are 10 ms apart. Red pacing marker illustrates location of pacing. **B.** Percent change in conduction velocity (vs. before light application) during and after application of light at different power during 500 ms CL pacing, for co-cultures of CMs with MFBs or ChR2-MFBs. Data for 10*I_0_ is not shown since almost all samples beat spontaneously with CL less than 500 ms at this intensity. * shows p<0.05, # shows p<0.005 difference between ChR2-MFB/CM and MFB/CM co-cultures.

Finally, light-induced inward current decreased action potential duration (APD), which reversed upon removal of light (Fig. 5A). Application of light to ChR2-MF/CM co-cultures resulted in a statistically significant decrease in APD at 80% repolarization (APD_80_) at 3*I_0_ in ChR2-MFB/CM co-cultures (−14±7% from a baseline of 161±11 ms, n=5, p=0.004, Fig. 5B). P-values for electrophysiological parameters calculated using other methods are provided in the Supplemental Table.

**Figure 5.**
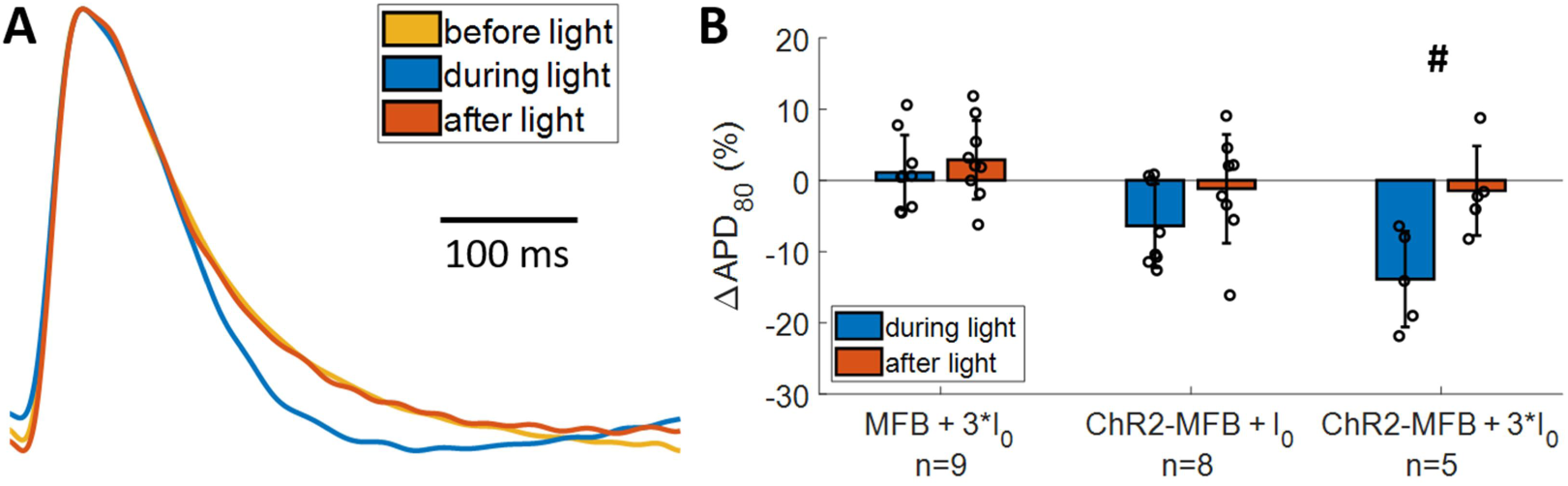
Inward currents in myofibroblasts change action potential morphology in co-cultured myofibroblast syncytia. **A.** Averaged AP trace before (gold), during (blue), and after (orange) application of 3*I_0_ blue light to activate ChR2 current in ChR2-MFBs. **B.** Percent change in action potential duration (vs. before light application) during and after application of light at different power during 500 ms CL pacing, for co-cultures of CMs with MFBs or ChR2-MFBs. * shows p<0.05, # shows p<0.005 difference between ChR2-MFB/CM and MFB/CM co-cultures.

Acute application of a very high level of light (1.2 mW/mm^2^) to ChR2-MFBs did not affect the MFBs’ contractility and force generation, as measured for single cells seeded on flexible micropost arrays (Supp. Fig. 2), excluding the possibility that inward currents in MFBs caused them to contract and potentially influence CMs by activating mechanosensitive channels.

### Insights from a mathematical model of the co-cultures of cardiomyocytes and ChR2-transduced myofibroblasts

To better understand these results, a numerical model of a line of 30 neonatal rat ventricular CMs connected to MFBs transduced with ChR2 currents was created (Fig. 1). Using maximum likelihood estimation, G_MFB-CM_ = 1 nS/CM and g_ChR2_ = 3.5 mS/μF were determined to best fit the experimental CL, ΔCV, and ΔAPD, with the model output falling within the 95% confidence interval for each parameter, and a geometric mean of p=0.17. To explore a broader parameter space, electrical connectivity between MFBs and CMs (G_MFB-CM_) and light intensity were varied. The model showed that at 3*I_0_ or G_MFB-CM_≤ 10^−0.5^ nS/CM, no spontaneous beating was produced, since either there was too little ChR2 current produced, or this current was unable to depolarize the CMs, respectively (Fig. 6A). However, at the G_MFB-CM_ determined by maximum likelihood estimation (1 nS/CM), spontaneous beating could exceed the pacing rate (500 ms CL) in response to light levels greater than between 3*I_0_ and 10*I_0_ (Fig. 6A, see blue dots). This behavior fit with our experiments, in which only 4/10 of our samples beat spontaneously at 3*I_0_, while 6/7 samples beat faster than the paced rate when stimulated with 10*I_0_. This higher level of light reduced CL by 22% in the model, similar to our experiments in which CL was reduced by 36±18% (n=7). Furthermore, the model predicted CMs could become inexcitable at high light levels and high G_MFB-CM_ (Fig. 6A, gray box), but only at G_MFB-CM_ greater than predicted and at light levels too high for us to apply while recording, due to optical crosstalk.

**Figure 6.**
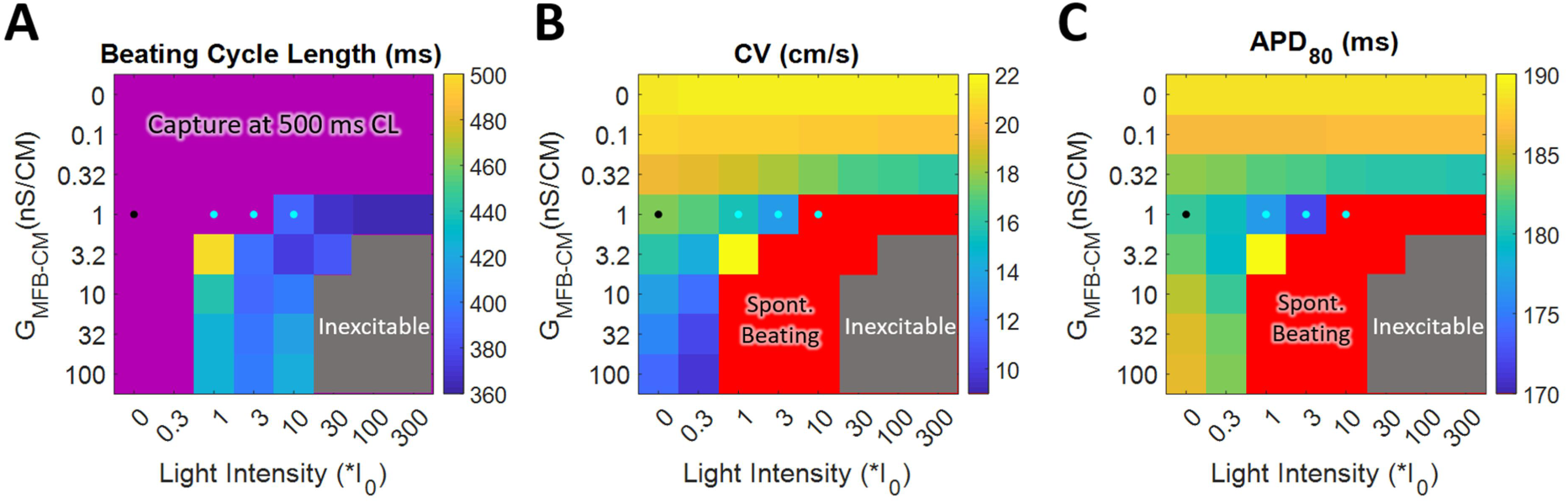
Linear cable model of MFBs on top of CMs in the same 1:2.5 cell ratio used in experiments verifies experimental findings. Beating cycle length (**A**), conduction velocity (**B**), and APD_80_ (**C**) for CMs paced at 500 ms CL at different MFB-CM conductance and light intensities. Purple region in A denotes spontaneous beating was slower than the paced CL. CV and APD_80_ were not calculated for the red region in B and C since spontaneous beating prevents 500 ms CL pacing. Gray region denotes that cells were inexcitable. Black dot denotes modeled values without light at the MFB-CM conductance with the maximum likelihood. Blue dots denote values at light levels from experiments at the MFB-CM conductance with the maximum likelihood.

The model also showed slowing with addition of MFBs, and this slowing increased with G_MFB-CM_ and light intensity (Fig. 6B). Similar to effects on spontaneous beating, if G_MFB-CM_ is low, inward currents from light application have very little effect on CMs, as expected since this current cannot pass to CMs. With G_MFB-CM_ of 1 nS/CM addition of MFBs alone decreased CV by 3.9 cm/s (Fig. 6B, compare black dot with top row, where G_MFB-CM_ = 0, which is equivalent to pure CM in the model). Application of I_0_ and 3*I_0_ light further decreased CV by 12% and 23%, respectively (Fig. 6B, compare blue dots with black dot), similar to experiments (Fig. 4B), while higher levels of light caused loss of capture due to spontaneous beating, as discussed in Fig. 6A.

Addition of MFBs to CMs decreased APD_80_ non-monotonically with increasing G_MFB-CM_, reducing APD by as much as 7 ms at 1 nS/CM, but by only 3 ms at 100 nS/CM (Fig. 6C, top row equivalent to the absence of MFBs). However, the effect of light on APD_80_ did increase monotonically with G_MFB-CM_, with APD_80_ decreasing by 3% and 5% for I_0_,and 3*I_0_ respectively, at G_MFB-CM_ = 1 nS/CM (Fig. 6C, compare blue dots with black dot), in line with experiments (Fig. 5B). Further investigation suggested that the decrease in APD in response to light observed experimentally (Fig. 5B) could be attributed to a more positive resting potential and decreased AP amplitude (Supp. Fig. 3A) instead of an increased repolarization rate, as would be suggested by a normalized trace (Supp. Fig 3B).

The model allowed further clarification of the mechanism of slowing, showing that co-culture with MFBs depolarized CMs during diastole from −71.8 to −67.8 mV (black dot for CM, Fig. 7A). Application of light produced additional inward ChR2 current that further depolarized CMs co-cultured with ChR2-MFBs to −65.0 or −62.5 mV for I_0_ or 3*I_0_, respectively (Fig. 7A). Depolarization of CMs decreased their inward sodium current from 169 pA/pF for pure CMs to 85 pA/pF for ChR2-MF/CM co-cultures, and to 50 and 18 pA/pF with the addition of I_0_ or 3*I_0_ light, respectively (Fig. 7B). This reduced inward current then resulted in slower CV (Fig. 7C). At 3*I_0_ (right-most blue dot in Fig. 7C), a significant portion of conduction (26%) was mediated by calcium current (compare right-most blue dots for I_Na_ and I_Ca_ in Fig. 7B). The magnitude of this calcium component was relatively unaffected by the changes in diastolic potential (Fig. 7B), leaving it available to support conduction and participate in spontaneous depolarization at higher diastolic potentials.

**Figure 7.**
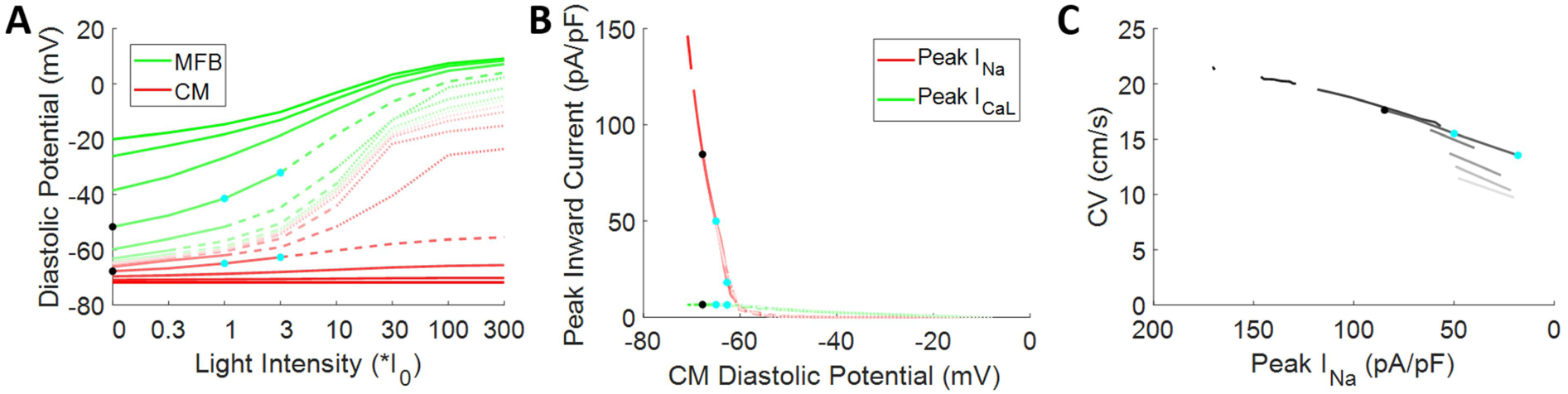
Modeling shows the process by which inward myofibroblast current slows cardiomyocyte conduction. **A.** Maximum diastolic potential of MFB (green) and CM (red) at different coupling levels and light intensities. **B.** Peak inward sodium current (red) and L-type calcium current (green) versus CM diastolic potential. **C.** Conduction velocity versus peak sodium current. MFB-CM coupling starts at 0 then varies by half log_10_ increments from 10^−1^ nS/CM (dark) to 10^2^ nS/CM (light). Solid lines show conditions that allowed capture at the 500 ms paced CL. Dashed lines show conditions that caused spontaneous beating. Dotted lines show conditions that caused cells to be inexcitable. Black dot denotes modeled values without light at the MF-CMB conductance with the maximum likelihood. Blue dots denote values at light levels from experiments at the MFB-CM conductance with the maximum likelihood.

## Discussion

Since the initial findings of Miragoli, et. al [16] that addition of MFBs to CM cultures causes RP elevation, conduction slowing, and spontaneous beating, a number of studies have attributed MFB-induced conduction slowing to electrical coupling between CMs and MFBs [14–18, 36]. Some studies used FRAP as evidence of diffusional (and therefore presumably electrical) coupling between CMs and MFBs [17], but these cannot be translated to a value of electrical conductance [37] or show that the connection is strong enough to cause significant slowing. Dual-cell patch clamp has been used to quantify the electrical connection between CM and MFB pairs [14, 38], but not in a syncytium, which allows for measurement of macroscopic CV, and has different electrophysiology than single cells [39]. Furthermore, dual-cell patch clamp measures conductance between cells next to each other, where due to the small height of cells relative to area only a small fraction of the cell surface is available for connection, although this is not the case for cells in 3-D tissue. In this study MFBs were seeded on top of CM monolayers, allowing MFBs to contact CMs over a large area and limiting their disruption of CM-CM connections. We also prioritized using both MFBs and CMs from the same source (neonatal rat ventricle) in our experiments, and building a computational model based on these same two cell types.

Interventional experiments have knocked down Cx43 in MFBs, and found that doing so increased CV in co-cultures, compared with CMs co-cultured with MFBs alone, to provide direct evidence of an electrical mechanism for CV reduction by MFBs [11, 18]. However, decreasing Cx43 expression also inhibits fibroblast differentiation to MFB [40, 41], so Cx43 knockdown alone could reduce the number of MFBs present and increase CV even if MFB-suppression of CV is by non-electrical means (e.g., paracrine or mechanical signaling). Studies that dynamically and specifically alter electrophysiology of cells putatively connected to CMs and monitor the subsequent changes in CM electrophysiology circumvent the confounding effects of Cx43-mediated changes in MFB differentiation. Previous studies have shown CM electrophysiology can be modulated by acutely altering exogenous potassium currents in co-cultured 3T3 fibroblasts [42, 43]. Another used 3T3 fibroblasts transduced with ChR2 and applied mW/mm^2^ light flashes to pace CMs [44]. Finally, a study used sphingosine-1-phosphate to increase MFB inward currents, and found that it suppressed CM excitability in co-cultures with MFBs, but not CMs alone [46].

In this study, light was used to produce steady inward current specifically in ChR2-MFBs. MFBs were sufficiently connected electrically to CMs for their coupling to produce ectopic beating (Fig. 3), conduction slowing (Fig. 4), and decreased APD_80_ (Fig. 5). The rapid time scale of these changes (within seconds) eliminates the possibility that changes in cardiac ion channel expression in response to the presence of MFBs underlie these effects. Additionally, the absence of changes in force generation with application of light to ChR2-MFBs (Supp. Fig. 2) rules out the possibility that CV slowing occurred secondary to acute changes in MFB tugging forces [12]. The fact that light had no effect on control MFB/CM co-cultures (Fig. 3-5) also supports the notion that the CV effects were due to light-induced ChR2 currents and not to off-target effects such as heating or photochemical reactions.

Quinn et al. [47] probed electrical connections between MFBs and CMs *in vivo* by creating genetically engineered mice that expressed an optogenetic voltage sensor specifically in non-CMs, and found time-varying signals specifically near a site of cryoinjury, suggesting that non-CMs can electrically connect to CMs in areas of injury. However, the promoter they used may not be limited to non-CMs, particularly in areas of injury [48]. Furthermore, non-CMs are better voltage followers than drivers when coupled to CMs because of their higher sarcolemmal resistance and lower sarcolemmal currents, as discussed in [37]. While the lack of an MFB-specific promoter remains a challenge, this study shows that using a similar design with an optogenetic actuator instead of an optogenetic sensor may be better suited to determine whether MFBs are sufficiently connected to CMs to cause conduction slowing and spontaneous beating *in vivo* under conditions of activated, inward MFB current. Other proposed mechanisms, such as MFBs being a current sink during the late phase of CM depolarization due to their outward currents and capacitance, remain to be explored. Additionally, paracrine and mechanical effects of MFBs may also contribute to slowing.

The effects of MFB addition to 2-D CM syncytia have been previously explored computationally [32, 49, 50]. However, only one previous study used a neonatal rat ventricular CM model [51]. It simulated a mixture of CMs and MFBs, allowing MFBs to interrupt CM-CM connections, and focused on how the resulting structural heterogeneity could produce wavebreaks and reentrant arrhythmia [51]. In this work, a modified neonatal rat ventricular CM model was used together with a MFB model parameterized directly from data from cultured MFBs [14] to examine how MFBs can affect macroscopic tissue electrophysiology in the absence of disrupted CM-CM connections.

In our experiments, cells were densely seeded with MFBs on top of CMs, so that the interface between MFBs and CMs occurred over a large area. Since G_MFB-CM_ is unknown in this configuration, and g_ChR2_ has not been measured in MFBs, a wide range of potential G_MFB-CM_ and values were investigated, and maximum likelihood estimation used to determine the values. The model replicates all of the phenomena seen in experiments, including spontaneous beating (Fig. 6A), conduction slowing (Fig. 6B), and APD reduction (Fig. 6C).

The model substantiates the previously proposed mechanism [11, 14, 16–18, 49, 50, 52] that inward currents in MFBs depolarize CMs, inactivating sodium channels, which slows conduction (Fig. 7). It also suggests that the sodium channel is largely inactivated when the CMs are depolarized (Fig. 7B), and conduction is mediated significantly by calcium currents, which are less affected by the resting potential (Fig. 7B). The block of sodium currents and significant calcium-mediated conduction after depolarization by MFBs is in agreement with a previous study that found little effect of the sodium channel blocker TTX on CV in CM strands co-cultured with large numbers of MFBs [16]. Additionally, the model explained why, even though ChR2 has a reversal potential near 10 mV and therefore might be expected to slow repolarization at voltages negative to 10 mV, application of light decreased APD_80_, both experimentally and in simulation (Fig. 5B and 6C). This occurred because it made the resting potential more positive (Fig. 7A) decreasing the amplitude of the action potential (Supp. Fig. 3A) and resulting in a shorter time to repolarize, given a similar repolarization rate (Supp. Fig. 3B). The fact that inward current reduces both AP amplitude and repolarization rate, each of which have opposite effects on APD, can explain the mixed effects that inward currents from MFBs have been found to have on APD_80_ experimentally [12, 17, 18, 53] and computationally [49, 50, 54–56].

## Conclusion

This study used optogenetic actuation of inward current in myofibroblasts to show they can acutely cause ectopic beating, conduction slowing, and decreased action potential duration in co-cultured cardiomyocytes, clearly demonstrating functional electrical coupling between myofibroblasts and cardiomyocytes in syncytium. With promoters specific for non-cardiomyocytes, this optogenetic actuation method could be used in vivo to conclusively demonstrate a functional connection between cardiomyocytes and myofibroblasts. Computational modeling of the experiments allowed us to further probe the process by which slowing occurs and suggests that the combination of cardiomyocyte/myofibroblast electrical coupling and myofibroblast currents reduce conduction velocity and produce spontaneous beating in cardiomyocytes, perhaps increasing the risk of life-threatening arrhythmia.

## Acknowledgements

This work was supported by NIH grant R01 HL127087. Special thanks to Dr. Gordon Tomaselli for discussion of the findings. Confocal imaging was supported by NIH grant S10 RR024550 (Dr. Scot Kuo). The authors would also like to acknowledge Aleksandra Klimas for expert advice on simultaneous optical recording and mapping, as well as Shoshana Das and Robert Hawthorne, who conducted experiments that were not used in the final version of this work.

