## Supplemental Figures for "Optogenetic currents in myofibroblasts acutely alter electrophysiology and conduction of co-cultured cardiomyocytes"

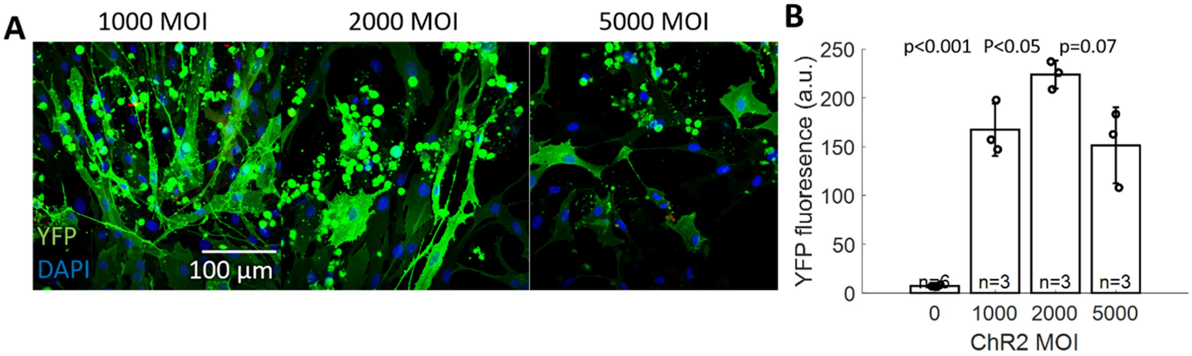


**Supplemental Figure 1. Transduction of myofibroblasts with ChR2. A.** Confocal images of MFBs transduced with varying multiplicity of infection (MOI) of ChR2 adenovirus. Green shows YFP, which marks cells transduced with ChR2, and blue shows DAPI, showing cell nuclei. Higher doses of virus were associated with some cell death (rounded cells). All images are taken at the same scale and settings. **B.** YFP fluorescence, as measured by a plate fluorescence reader, was significantly increased by transducing with 2000 MOI versus 1000 MOI, while 5000 MOI decreased YFP fluorescence. Therefore, MFBs were transduced at approximately 2000 MOI in future experiments.

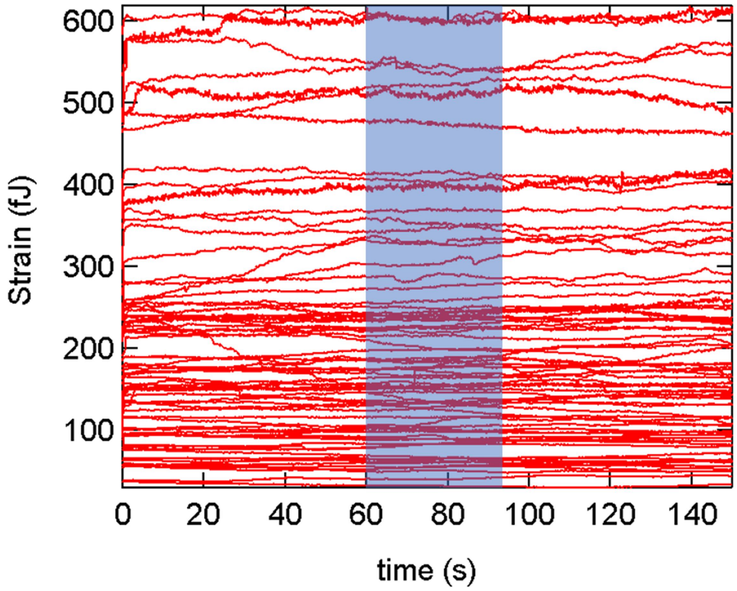


**Supplemental Figure 2. ChR2-transduced myofibroblast strain energy during excitation by light**. Recordings of ChR2-MFB strain energy before, during (blue box), and after excitation with 1.2 $mW/mm^2$ blue light shows that opening of ChR2 channels does not acutely affect contractile energy.

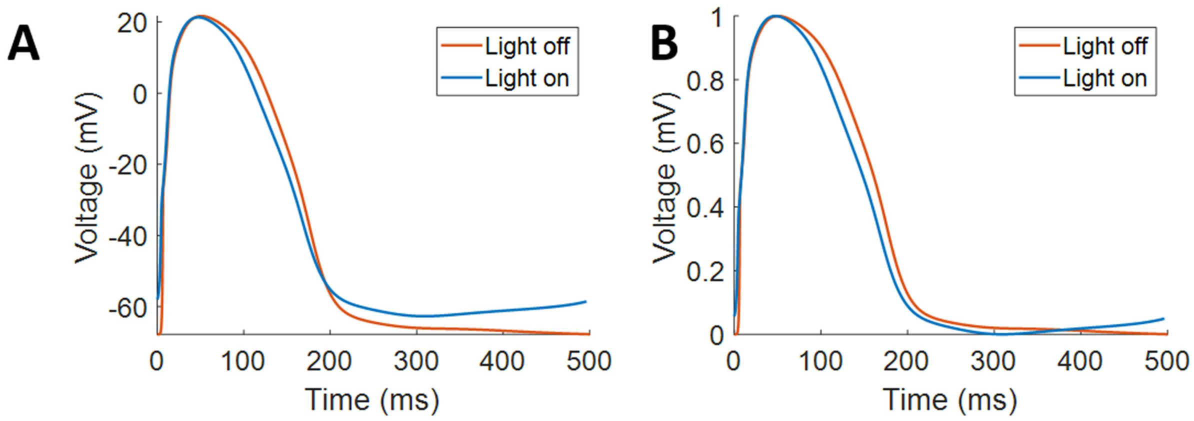


**Supplemental Figure 3. Amplitude changes contribute to changes in action potential duration.**
**A.** Modelled AP traces without (blue) and with (orange) $3*I_0$ light. **B.** Same as A, but with normalized AP traces, as are measured during optical mapping.

| **A** | Paired p ($\Delta_{light\ on}$) | | Paired p ($\Delta_{light\ off}$) | | $\Delta_{light\ on}$ p(ChR2 vs. no ChR2) | | 95% CI of during light (CL) or after light-before light | | | |
|---|---|---|---|---|---|---|---|---|---|---|
| | Low | High | Low | High | Low | High | Low | | High | |
| CL | 0.081 | 0.002 | 0.104 | 0.002 | 0.079 | 0.002 | 499 | 501 | 499 | 500 |
| CV | 0.025 | 0.005 | 0.013 | 0.009 | 0.017 | 0.003 | -1.8 | 1.7 | -0.6 | 0.07 |
| $APD_{80}$ | 0.019 | 0.024 | 0.078 | 0.007 | 0.895 | 0.013 | -11 | 8 | -15 | 8 |

| **B** | Paired p (%$\Delta_{light\ on}$) | | Paired p (%$\Delta_{light\ off}$) | | %$\Delta_{light\ on}$ p(ChR2 vs. no ChR2) | | 95% CI of %$\Delta_{after\ light-before\ light}$ | | | |
|---|---|---|---|---|---|---|---|---|---|---|
| | Low | High | Low | High | Low | High | Low | | High | |
| CL | 0.081 | 0.002 | 0.104 | 0.002 | 0.076 | 0.002 | -9.2 | 9.3 | -0.3 | 0.1 |
| CV | 0.018 | $<10^{-3}$ | 0.010 | $<10^{-3}$ | 0.011 | $<10^{-3}$ | -12.6 | 11.0 | -5.1 | 0.7 |
| $APD_{80}$ | 0.018 | 0.010 | 0.083 | 0.003 | 0.678 | 0.004 | -7.5 | 5.2 | -9.2 | 6.4 |

**Supplemental Table. Additional p-values and confidence intervals.** For absolute (**A**) and relative (**B**) difference for measured parameters for low ($3*I_0$ for CL, and $I_0$ for others) and high ($10*I_0$ for CL and $3*I_0$ for others) light levels. $p<0.05$ shaded.
